# Science with no fiction: measuring the veracity of scientific reports by citation analysis

**DOI:** 10.1101/172940

**Authors:** Peter Grabitz, Yuri Lazebnik, Josh Nicholson, Sean Rife

## Abstract

The current crisis of veracity in biomedical research is enabled by the lack of publicly accessible information on whether the reported scientific claims are valid. One approach to solve this problem is to replicate previous studies by specialized reproducibility centers. However, this approach is costly or unaffordable and raises a number of yet to be resolved concerns that question its effectiveness and validity. We propose to use an approach that yields a simple numerical measure of veracity, the R-factor, by summarizing the outcomes of already published studies that have attempted to test a claim. The R-factor of an investigator, a journal, or an institution would be the average of the R-factors of the claims they reported. We illustrate this approach using three studies recently tested by a replication initiative, compare the results, and discuss how using the R-factor can help improve the veracity of scientific research.

---

The current crisis of veracity in biomedical research, and having less than a half of preclinical studies reproducible (Begley & Ellis, 2012; Prinz, Schlange, & Asadullah, 2011) is truly a crisis, has spilled from a discussion in scientific journals (Begley & Ellis, 2012; Casadevall & Fang, 2010; Collins & Tabak, 2014; Fang, Steen, & Casadevall, 2012; Freedman, Cockburn, & Simcoe, 2015; Ioannidis, 2005, 2017; Leek & Jager, 2017) into the pages of national newspapers (Angell, 2009; Carey, 2015; Glanz, 2017) and popular books with provocative titles (Harris, 2017). This development suggests that scientists might need to put their house in order before asking for more money to expand it.

The approaches that have been tried or proposed are: calling on scientists to be better and “publish houses of brick, not mansions of straw” (Kaelin, 2017), perhaps under the scrutiny of video surveillance in the laboratory (Clark, 2017), requiring raw data and additional information when submitting an article (Editorial, 2017a) or a funding report (https://grants.nih.gov/reproducibility/index.htm), and establishing reproducibility initiatives that replicate prior studies to serve as a deterrent for future abuse of scientific rigor. One of these initiatives, Reproducibility Project: Cancer Biology, was organized following the report that only 6 out of 53 landmark cancer research studies could be verified (Begley & Ellis, 2012) and set to replicate 50 cancer research reports out of 290,444 published by the field between 2010 and 2012 (Errington et al., 2014). The reports on replicating the first seven studies have been published this year (Aird, Kandela, Mantis, & Reproducibility Project: Cancer, 2017; Horrigan et al., 2017; Horrigan & Reproducibility Project: Cancer, 2017; Kandela, Aird, & Reproducibility Project: Cancer, 2017; Mantis, Kandela, Aird, & Reproducibility Project: Cancer, 2017; Shan, Fung, Kosaka, Danet-Desnoyers, & Reproducibility Project: Cancer, 2017; Showalter et al., 2017).

We would like to use these reports to suggest how the credibility crisis can be solved effectively and at a relatively small cost by assigning each published scientific claim a simple measure of veracity, which we call the R-factor (Nicholson & Lazebnik, 2014), with R standing for reproducibility, reputation, responsibility, and robustness.

The R-factor is based on the same rule of science that underlies the replication initiatives, namely that a scientific claim should be independently confirmed before accepting it as fact. Hence, it is calculated simply by dividing the number of published reports that have verified a scientific claim by the number of attempts to do so; to emphasize, this calculation excludes citations that merely mention the claim without testing it. The result is a number between 0 (claim is unverified) and 1 (claim is confirmed). The R-factor of an investigator, a journal, or an institution would be the average of the R-factors of the claims they reported.

The R-factor is also based on another principle of science, that new research should proceed from a comprehensive understanding of previous work. Following this principle is becoming more difficult because the sheer number of publications overwhelms even experts, thus making their expertise even more narrow. The R-factor would help to solve this problem not only by providing a measure of veracity for a published claim, but also by indicating the studies that verified or refuted it.

Let us illustrate the approach we propose using three of the cases evaluated by the reproducibility project and then discuss how using the R-factor can help make biomedical research more trustworthy.

## Cases

### Case 1: The Common Feature of Leukemia-Associated IDH1 and IDH2 Mutations Is a Neomorphic Enzyme Activity Converting a-Ketoglutarate to 2-Hydroxyglutarate (Ward et al., 2010)

IDH1 and IDH2 are metabolic enzymes that are often mutated in certain cancers. The report by Ward et al. claims, as its title indicates, that all these mutants make 2-hydroxyglutarate, a molecule that is not produced by the normal enzymes and which is present in normal cells in trace amounts. The significance of this claim is in explaining how distinct IDH1 and IDH2 mutations contribute to cancer development.

By reviewing 743 articles citing the report by Ward et al., we identified 17 independent studies, (Table S1), including the replication report (Showalter et al., 2017), which verified the claim by measuring the activities of mutants IDH1 and IDH2. We considered a study independent if its senior author was different from that of the cited report, which included collaborative studies with the laboratory reporting the claim. One out of 17 confirming articles was collaborative in this case. We have found no refuting studies. Hence, the R-factor of the claim by Ward et al. is 17/(17+0) = 1.0_17_ (Figure 1, left), which is concordant with the conclusion of the replication study (Showalter et al., 2017).

**Figure 1.**
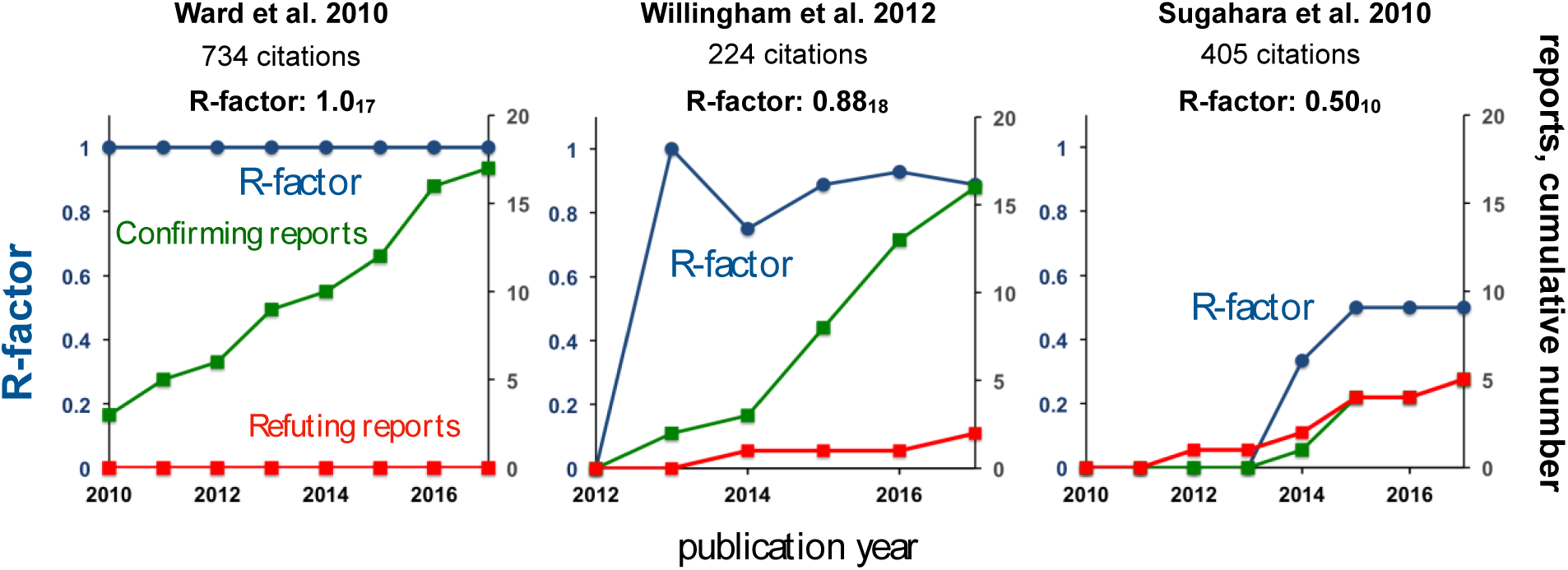
The R-graphs of three replication studies. The articles citing the three reports were found by searching the Web of Science and reviewed to identify the studies that confirmed or refuted the claims of the reports. The R-factor was calculated by dividing the number of confirming studies on the sum of confirming and refuting studies, which is indicated in subscript.

### Case 2: The CD47-signal regulatory protein alpha (SIRPα) interaction is a therapeutic target for human solid tumors (Willingham et al., 2012)

CD47 is a ubiquitously expressed membrane protein with many functions, one of which is to signal “do not eat me” to macrophages and dendritic cells by binding SIRPα, a protein present on their surface (Matlung, Szilagyi, Barclay, & van den Berg, 2017). Willingham et al. have reported that antibodies to CD47 that disrupt the CD47-SIRPα interaction inhibit the growth and metastasis of solid human tumors explanted into mice, and thus claimed that this interaction is a therapeutic target for human solid tumors.

The replication report (Horrigan & Reproducibility Project: Cancer, 2017) refuted the claim by finding no statistically significant effect of a CD47 antibody on tumor growth in an animal model, with the stated caveats that some of the tumors regressed spontaneously in the control group, that only one of the CD47 antibodies used in the original study was used in the replication study, and that only one out of several models used in the original study was used to test its claim.

By reviewing 224 reports citing the study by Willingham et al., we have identified 16 independent studies (Table S1) that confirmed the claim. These studies used several CD47 antibodies, SIRPα mimetics, and other approaches. Two citing studies, including the replication report, refuted the claim (Horrigan & Reproducibility Project: Cancer, 2017; Soto-Pantoja et al., 2014). Hence, the R-factor of the claim is 16/(16+2) = 0.88_18_ (Figure 1, middle), which is discordant with the conclusion of the replication study but concordant with the current standing of the claim in the field, the debate about the complexity of the underlying mechanisms notwithstanding (Matlung et al., 2017).

### Case 3: Coadministration of a tumor-penetrating peptide enhances the efficacy of cancer drugs (Sugahara et al., 2010)

The authors of this report previously found that conjugating the peptide iRGD to various cancer drugs makes these drugs more potent by helping them penetrate into target tissues (Sugahara et al., 2009). In this study, Sugahara et al. claim that the same effect can be achieved by injecting free iRGD together with a cancer drug, thus bypassing the need to link the peptide chemically.

The replication study found no evidence supporting the claim (Mantis, Kandela, Aird, & Reproducibility Project Canc, 2017).

By reviewing 405 citing publications, we have identified nine studies that supported the claim and five that refuted it (Table S1). Two of the studies (Akashi et al., 2014; Sugahara et al., 2015) were included in both categories, as some of the reported experiments supported the claim while others did not, apparently because the effect depended on the experimental system used.

We then excluded two of supporting reports because although they tested different drugs, the graph of tumor growth presented in the first report (Q. Zhang et al., 2015) was superimposable, except of the labeling, to the graph in the second (Y. Zhang et al., 2016). In two other supporting studies the graphs representing the survival of treated mice implied that the animals died surprisingly regularly, one mouse per each observation point in each of the thirteen cohorts (six groups of six mice each (Gu et al., 2013), and seven groups of 10 mice (Wang et al., 2014)). Because we considered such orderly demise improbable and an expert in animal experiments concurred with our opinion, we excluded these studies from analysis.

The remaining five confirming and five refuting studies resulted in the R-factor = 5/(5+5) = 0.50_10_ (Figure 1, right).

### What are the benefits of calculating the R-factor?

Perhaps the best way to begin answering this question is by mentioning a recent study (Benjamin, Mandel, & Kimmelman, 2017) in which 196 cancer researchers, including 138 experts, were asked to predict whether the reports the reproducibility project was set to verify, including two studies we have analyzed (Sugahara et al., 2010; Willingham et al., 2012), are reproducible. The authors concluded that the scientists were poor forecasters who overestimated the validity of the studies (Benjamin et al., 2017).

We would like to suggest that the scientists would do much better if they could see the R-graphs of the studies in question (Fig. 1). For example, knowing that 8 out of 9 studies that tested a claim confirmed it (Fig. 1, middle, year 2015, when the study by Benjamin et al. began) would not only make the prediction more accurate, if not easy, but would also raise the question whether the tenth attempt to verify this claim is justified and, once the replication study found that the claim is irreproducible, whether this conclusion itself needs an independent review. Instead, the scientists outside of the narrow field had to rely on their intuition because the required information was not readily available. This is the deficiency that calculating the R-factor and making the results freely available can correct.

The R-factor is relatively easy to calculate, as the process requires no laboratory equipment, laboratory animals, or reagents, and can be done by anyone with a general expertise in biomedical research. This calculation is also much faster than experimental replication: all three studies (Fig.1) were evaluated during one week by one person.

Since the R-factor uses not one, but all reports that have evaluated a claim (10 to 18 in the examples we used), one can argue that the confidence level that the R-factor provides is at least as valid as that provided by a replication study, unless no reports citing the claim of interest are available, in which case a replication study is in order.

The R-factor is universal in that it is applicable to any scientific claim, based on either experimental or theoretical work, and, by extension, to individual researchers, laboratories, institutions, countries, or any other group, with no basic constraints on how many reports produced by these groups can be evaluated. This feature implies that the R-factor can be calculated for each claim made in a report, should it make more than one.

Since the R-factor can be anywhere between 0 and 1, it reflects the realities of experimental science, where a binary scale of right and wrong is not always applicable, especially at the initial stages of developing an idea, or when the complexity of the experimental system calls for time to find the final answer. For example, the R-factor of 1.0 for the claim by Ward et al. can be explained by the fact that the claim can be verified unambiguously by measuring activity of the IDH mutants with an established approach. The R-factor of 0.88 for the claim by Willingham et al. may reflect the debate on whether the mechanisms underlying the effect of CD47 antibodies are more complex than initially envisioned (reviewed in (Matlung et al., 2017)).

The R-factor of 0.5 for the claim by Sugahara et al. gives a warning that the claim might be untrue, which may be a surprise for the reader who relies on the citation indexes and impact factors, as the article has been cited 405 times and has been published in *Science*, a top journal. However, the R-factor of 0.5 also leaves open the possibility that the claimed approach is applicable to some systems and suggests that further testing is needed, which is where the replication initiatives can be very helpful. The cases like that of Sugahara et al. and the opportunity to contribute to evaluating them through the R-factor might invite researchers to report unsuccessful attempts to rest reported claims, as so called negative results often go unpublished because they are considered inconsequential.

Because the R-factor relies on experimental reports from experts in the field, this approach alleviates or bypasses the concerns associated with replication initiatives (Bissell, 2013; Editorial, 2017b), such as lack of technical expertise or of suitable experimental models in a laboratory specialized in replicating prior studies. This approach also bypasses the debate on what it means to replicate a study, as it merely asks whether the main claim of a study, typically formulated in the title of the report, is confirmed or not. For example, the ongoing clinical trials of CD47 antibodies (Matlung et al., 2017) cannot in principle replicate the study by Willingham et al., as it used mice, but the trials would confirm or refute its main claim.

Finally, the R-factor and the information that comes with it (Fig. 1, Table 1S) allows a researcher to focus on the articles that tested the claim, the opportunity that can be especially valuable for the highly cited reports, as the majority of these citations (97.7%, 92%, and 97.5% for cases 1-3) merely mention the cited report without evaluating it experimentally. As previous studies have illustrated (Greenberg, 2009), the sheer number of mentioning citations, especially if aided by their skillful use, can make a field accept a dubious claim as a fact.

### How can the R-factor help to solve the crisis of credibility?

Assigning the R-factor to scientific reports will solve two interrelated and unresolved problems that underlie the crisis: that the careers of researchers are disconnected from the veracity of what they report, and that finding the veracity of a claim or a researcher is currently difficult.

The current choices to determine whether a scientific claim or a scientist is reliable are to consult insiders in the field, which may require certain connections to have a frank assessment and presumes that the insiders are not misled themselves, to review dozens to thousands of articles that cite the report of interest, or to replicate the study independently, which could be expensive or, at the times of financial constraints, unaffordable.

With the absence of easily accessible information and transparency about the reliability of reported claims, and the deluge of publications that can overwhelm even an expert, the careers of academic researchers are affected little by the veracity of what they publish or the lack thereof, short of scandals associated with outright fraud, but instead depend on the number of published articles, the number of citations, and the impact factors (citation indexes) of the journals (Fig. 2, left).

**Figure 2.**
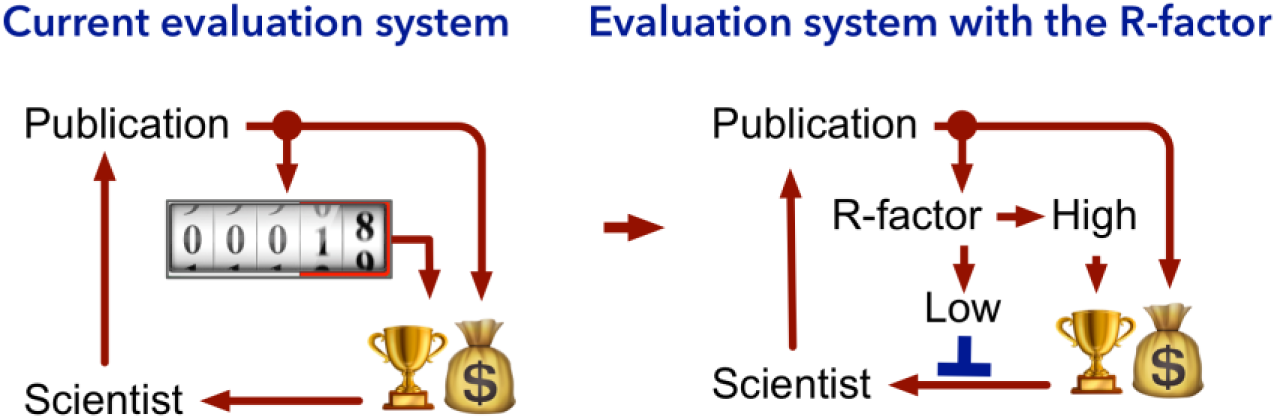
The difference using the R-factor can make. **Left**: The pressure to publish more articles is unrestrained by the lack of veracity in what is published. The counter in the center reflects the fact that what currently matters for receiving the rewards, that is salaries, research funding, and recognition are the number of articles a scientist publishes, the number of citations they receive, and the impact factor (the average number of citations per article) of the journal in which they appear. **Right**: Introducing the R-factor, which reports whether a claim has been verified, inhibits the rush to publish unverified claims (blue line) by linking the veracity of publications to life and research support, thus correcting the existing imbalance of incentives.

Having the R-factor indicated on the first page of the report, in much the same way as the Altmetrics logo now informs the reader how popular the study is at social networks (Warren, Raison, & Dasgupta, 2017), would give anyone a numerical estimate of the report’s trustworthiness. Having this information openly and freely available, used, and discussed would enable, perhaps even force the evaluation system to consider veracity of the reports and the investigator in decisions related to their career (Fig. 2, right), and give the public a tool to judge for themselves. Importantly for the fairness of these decisions and judgments, the R-factor would reflect not a single replication study, but the sum of all reported attempts to reproduce the claim. Likewise, the credibility of an investigator would be estimated by the average veracity of all claims that they have reported, not unlike the batting average in baseball.

Because the R-factor can change over time (Fig. 1), and, in contrast to citation indexes that cannot decrease, not always to the better, our approach can help to change the current perception that a publication is a trophy, which, once acquired, would shine in the resume of its author forever supported by the citation index of the journal in which the report appeared. Instead, the worry that the R-factor can change to the worse for everyone to see is likely to make the authors, especially those who do the experiments but sometimes have little say on how the results are represented in the publication, more vigorous in insisting that the data and the conclusions are verified before submitting the manuscript.

Of course, the R-factor has its share of shortcomings and by no means an ideal measure of scientific excellence and not a panacea by itself. However, as Churchill said about democracy, “[it] is the worst form of Government except for all those other forms that have been tried from time to time.” We suggest the same can be said about the principle that scientific claims must be independently verified before accepting them as facts. The R-factor would help to apply and represent this principle in an easy to understand and easy to use form (Fig. 1), providing a sorely missed feedback in the evaluation system that governs biomedical research.

Although calculating the R-factor for a handful of reports is relatively simple, especially to an expert in the field, the question is who will calculate the R-factors for the thousands of researchers and their hundreds of thousands or even millions of reports. While these numbers look overwhelming, they are finite. We suggest that they can be processed using two complementary approaches – the collaboration of scientists who can calculate the R-factors for each other’s studies, and the application of machine learning technology, which brought such marvels as automatic language translation and face recognition from science fiction stories into our smartphones and has made great advances in analyzing the meaning of texts (Westergaard, Stærfeldt, Tønsberg, Jensen, & Brunak, 2017). A field that has elicited more credibility concerns than others, with cancer research being a primary candidate, could be a place to start.

Introducing the R-factor will be disruptive as it will bruise some egos and will disrupt the comfort of some scientific administrators. We feel, however, that this disruption will benefit future patients by giving a career advantage to the creative researchers and administrators who are committed to making biomedical research more productive. This change will help to restore public trust in science, which is now trending in the wrong direction.

We invite you to calculate the R-factor for the articles you like, dislike, or the articles that puzzle you. If you would like, you can also calculate your R-factor. What is it?

## Competing interests

The authors are co-founders of Verum Analytics, a company that has been created to make the R-factor of scientific claims freely and openly accessible and widely used.

## REFERENCES

Aird, F., Kandela, I., Mantis, C., & Reproducibility Project: Cancer, Biology. (2017). Replication Study: BET bromodomain inhibition as a therapeutic strategy to target c-Myc. Elife, 6. doi: 10.7554/eLife.21253

Akashi, Y., Oda, T., Ohara, Y., Miyamoto, R., Kurokawa, T., Hashimoto, S., … Ohkohchi, N. (2014). Anticancer effects of gemcitabine are enhanced by co-administered iRGD peptide in murine pancreatic cancer models that overexpressed neuropilin-1. British Journal of Cancer, 110(6), 1481–1487. doi: 10.1038/bjc.2014.49

Angell, Marcia. (2009). Drug companies and doctors: A story of curruption. The New York Review of Books. http://www.nybooks.com/articles/2009/01/15/drug-companies-doctorsa-story-of-corruption/

Begley, C. G., & Ellis, L. M. (2012). Drug development: Raise standards for preclinical cancer research. Nature, 483(7391), 531–533. doi: 10.1038/483531a

Benjamin, D., Mandel, D. R., & Kimmelman, J. (2017). Can cancer researchers accurately judge whether preclinical reports will reproduce? PLoS Biol, 15(6), e2002212. doi: 10.1371/journal.pbio.2002212

Bissell, M. (2013). Reproducibility: The risks of the replication drive. Nature, 503(7476), 333–334. doi:10.1038/503333a

Carey, B. (2015). Many Psychology Findings Not as Strong as Claimed, Study Says, New York Times. http://www.nytimes.com/2015/08/28/science/many-social-science-findings-not-as-strong-as-claimed-study-says.html

Casadevall, A., & Fang, F. C. (2010). Reproducible science. Infect Immun, 78(12), 4972–4975. doi: 10.1128/IAI.00908-10

Clark, T. D. (2017). Science, lies and video-taped experiments. Nature, 542(7640), 139. doi: 10.1038/542139a

Collins, F. S., & Tabak, L. A. (2014). Policy: NIH plans to enhance reproducibility. Nature, 505(7485), 612–613. doi: 10.1038/505612a.

Editorial. (2017a). Announcement: Transparency upgrade for Nature journals. Nature, 543(7645), 288. doi: 10.1038/543288b

Editorial. (2017b). The challenges of replication. Elife, 6., Jan 2017. doi: 10.7554/eLife.23693

Errington, T. M., Iorns, E., Gunn, W., Tan, F. E., Lomax, J., & Nosek, B. A. (2014). An open investigation of the reproducibility of cancer biology research. Elife, 3. doi: 10.7554/eLife.04333

Fang, F. C., Steen, R. G., & Casadevall, A. (2012). Misconduct accounts for the majority of retracted scientific publications. Proc Natl Acad Sci U S A, 109(42), 17028–17033. doi: 10.1073/pnas.1212247109

Freedman, L. P., Cockburn, I. M., & Simcoe, T. S. (2015). The Economics of Reproducibility in Preclinical Research. PLoS Biol, 13(6), e1002165. doi: 10.1371/journal.pbio.1002165

Glanz, J., Armendariz, A. (2017). Years of Ethics Charges, but Star Cancer Researcher Gets a Pass, New York Times. Retrieved from http://www.nytimes.com/2017/03/08/science/cancer-carlo-croce.html

Greenberg, S. A. (2009). How citation distortions create unfounded authority: analysis of a citation network. BMJ, 339, b2680. doi: 10.1136/bmj.b2680

Gu, G. Z., Gao, X. L., Hu, Q. Y., Kang, T., Liu, Z. Y., Jiang, M. Y., … Chen, J. (2013). The influence of the penetrating peptide iRGD on the effect of paclitaxel-loaded MT1-AF7p-conjugated nanoparticles on glioma cells. Biomaterials, 34(21), 5138–5148. doi: 10.1016/j.biomaterials.2013.03.036

Harris, Richard. (2017). Rigor Mortis. How sloppy science creates worthless cures, crushes hope, and wastes billions. New York: Basic Books.

Horrigan, S. K., Courville, P., Sampey, D., Zhou, F., Cai, S., & Reproducibility Project: Cancer, Biology. (2017). Replication Study: Melanoma genome sequencing reveals frequent PREX2 mutations. Elife, 6. doi: 10.7554/eLife.21634

Horrigan, S. K., & Reproducibility Project: Cancer, Biology. (2017). Replication Study: The CD47-signal regulatory protein alpha (SIRPa) interaction is a therapeutic target for human solid tumors. Elife, 6. doi: 10.7554/eLife.18173

Ioannidis, J. P. (2005). Why most published research findings are false. PLoS Med, 2(8), e124. doi: 10.1371/journal.pmed.0020124

Ioannidis, J. P. (2017). Acknowledging and Overcoming Nonreproducibility in Basic and Preclinical Research. JAMA, 317(10), 1019–1020. doi: 10.1001/jama.2017.0549

Kaelin, W. G., Jr. (2017). Publish houses of brick, not mansions of straw. Nature, 545(7655), 387. doi: 10.1038/545387a

Kandela, I., Aird, F., & Reproducibility Project: Cancer, Biology. (2017). Replication Study: Discovery and preclinical validation of drug indications using compendia of public gene expression data. Elife, 6. doi: 10.7554/eLife.17044

Leek, Jeffrey T., & Jager, Leah R. (2017). Is Most Published Research Really False? Annual Review of Statistics and Its Application, 4(1), 109–122. doi: 10.1146/annurev-statistics-060116-054104

Mantis, C., Kandela, I., Aird, F., & Reproducibility Project Canc, Biol. (2017). Replication Study: Coadministration of a tumor-penetrating peptide enhances the efficacy of cancer drugs. Elife, 6. doi: 10.7554/eLife.17584

Mantis, C., Kandela, I., Aird, F., & Reproducibility Project: Cancer, Biology. (2017). Replication Study: Coadministration of a tumor-penetrating peptide enhances the efficacy of cancer drugs. Elife, 6. doi: 10.7554/eLife.17584

Matlung, H. L., Szilagyi, K., Barclay, N. A., & van den Berg, T. K. (2017). The CD47-SIRPalpha signaling axis as an innate immune checkpoint in cancer. Immunol Rev, 276(1), 145–164. doi: 10.1111/imr.12527

Nicholson, Joshua, & Lazebnik, Yuri. (2014). The R-Factor: A Measure of Scientific Veracity. The Winnower. doi: 10.15200/winn.140832.20404

Prinz, F., Schlange, T., & Asadullah, K. (2011). Believe it or not: how much can we rely on published data on potential drug targets? Nat Rev Drug Discov, 10(9), 712. doi: 10.1038/nrd3439-c1

Shan, X., Fung, J. J., Kosaka, A., Danet-Desnoyers, G., & Reproducibility Project: Cancer, Biology. (2017). Replication Study: Inhibition of BET recruitment to chromatin as an effective treatment for MLL-fusion leukaemia. Elife, 6. doi: 10.7554/eLife.25306

Showalter, M. R., Hatakeyama, J., Cajka, T., VanderVorst, K., Carraway, K. L., Fiehn, O., & Reproducibility Project: Cancer, Biology. (2017). Replication Study: The common feature of leukemia-associated IDH1 and IDH2 mutations is a neomorphic enzyme activity converting alpha-ketoglutarate to 2-hydroxyglutarate. Elife, 6. doi: 10.7554/eLife.26030

Soto-Pantoja, D. R., Terabe, M., Ghosh, A., Ridnour, L. A., DeGraff, W. G., Wink, D. A., … Roberts, D. D. (2014). CD47 in the Tumor Microenvironment Limits Cooperation between Antitumor T-cell Immunity and Radiotherapy. Cancer Research, 74(23), 6771–6783. doi: 10.1158/0008-5472.an-14-0037-t

Sugahara, K. N., Scodeller, P., Braun, G. B., de Mendoza, T. H., Yamazaki, C. M., Kluger, M. D., … Lowy, A. (2015). A tumor-penetrating peptide enhances circulation-independent targeting of peritoneal carcinomatosis. Journal of Controlled Release, 212, 59–69. doi: 10.1016/j.jconrel.2015.06.009

Sugahara, K. N., Teesalu, T., Karmali, P. P., Kotamraju, V. R., Agemy, L., Girard, O. M., … Ruoslahti, E. (2009). Tissue-penetrating delivery of compounds and nanoparticles into tumors. Cancer Cell, 16(6), 510–520. doi: 10.1016/j.ccr.2009.10.013

Sugahara, K. N., Teesalu, T., Karmali, P. P., Kotamraju, V. R., Agemy, L., Greenwald, D. R., & Ruoslahti, E. (2010). Coadministration of a tumor-penetrating peptide enhances the efficacy of cancer drugs. Science, 328(5981), 1031–1035. doi: 10.1126/science.1183057

Wang, K., Zhang, X. F., Liu, Y., Liu, C., Jiang, B. H., & Jiang, Y. Y. (2014). Tumor penetrability and anti-angiogenesis using iRGD-mediated delivery of doxorubicin-polymer conjugates. Biomaterials, 35(30), 8735–8747. doi: 10.1016/j.biomaterials.2014.06.042

Ward, P. S., Patel, J., Wise, D. R., Abdel-Wahab, O., Bennett, B. D., Coller, H. A., … Thompson, C. B. (2010). The common feature of leukemia-associated IDH1 and IDH2 mutations is a neomorphic enzyme activity converting alpha-ketoglutarate to 2-hydroxyglutarate. Cancer Cell, 17(3), 225–234. doi: 10.1016/j.ccr.2010.01.020

Warren, H. R., Raison, N., & Dasgupta, P. (2017). The Rise of Altmetrics. JAMA, 317(2), 131–132. doi: 10.1001/jama.2016.18346

Westergaard, David, Stærfeldt, Hans-Henrik, Tønsberg, Christian, Jensen, Lars Juhl, & Brunak, Søren. (2017). Text mining of 15 million full-text scientific articles. bioRxiv. https://doi.org/10.1101/162099

Willingham, S. B., Volkmer, J. P., Gentles, A. J., Sahoo, D., Dalerba, P., Mitra, S. S., … Weissman, I. L. (2012). The CD47-signal regulatory protein alpha (SIRPa) interaction is a therapeutic target for human solid tumors. Proc Natl Acad Sci U S A, 109(17), 6662–6667. doi: 10.1073/pnas.1121623109

Zhang, Q., Zhang, Y., Li, K., Wang, H. Y., Li, H. Z., & Zheng, J. N. (2015). A Novel Strategy to Improve the Therapeutic Efficacy of Gemcitabine for Non-Small Cell Lung Cancer by the Tumor-Penetrating Peptide iRGD. PLoS One, 10(6). doi: 10.1371/journal.pone.0129865

Zhang, Y., Yang, J., Ding, M. H., Li, L. T., Lu, Z., Zhang, Q., & Zheng, J. N. (2016). Tumor-penetration and antitumor efficacy of cetuximab are enhanced by co-administered iRGD in a murine model of human NSCLC. Oncology Letters, 12(5), 3241–3249. doi: 10.3892/ol.2016.5081

